# *Absence of Evidence is Not Evidence of Absence:* The many flaws in the case against transgenerational epigenetic inheritance of pathogen avoidance in *C. elegans*

**DOI:** 10.1101/2024.06.07.597568

**Authors:** Rachel Kaletsky, Rebecca Moore, Titas Sengupta, Renee Seto, Borja Ceballos-Llera, Coleen T. Murphy

## Abstract

After examining the data and methods presented in Gainey, et al., ***bioRxiv***, 2024^1^ we conclude that the authors did not use an experimental paradigm that would have allowed them to replicate our results on transgenerational epigenetic inheritance (TEI) of learned avoidance that we regularly observe^2–5^. That is, we agree with the authors that their experiments show no evidence of TEI. However, there are substantial differences in their execution of every step of the work that also make it impossible for the authors to claim that they are replicating our experiments or protocols. Based on these differences, we do not believe there is an issue of “robustness” and “reliability” in our TEI findings, but rather that Hunter and colleagues have in fact not tested the *central* condition of TEI - that is, small RNA production by bacteria and subsequent uptake by *C. elegans* - nor have they carried out proper behavioral and imaging assays to assess this behavior. Our subsequent work shows that indeed this example of transgenerational epigenetic inheritance is not just observed in laboratory settings with PA14, but is also induced by wild strains of *Pseudomonas*, exhibiting its robustness. Just as we offered advice and training to Hunter and colleagues, we are happy to advise anyone who wishes to learn this assay.

We have now performed the experiments in the absence of azide, a deviation from our protocol that Hunter and colleagues deliberately made, and found that this omission may account for most if not all of the differences from our results. It is disingenuous for the authors to have presented their work as if they have used our protocol, given the fact that they chose to not use the same conditions for the most basic assay used in the work, the chemotaxis assay, which would have been necessary to replicate in order to reproduce our work. Therefore, Gainey et al.’s claims that our protocol or results are “irreproducible” are not supported by their evidence.

## Introduction

The opportunistic human pathogen PA14 “switches on” the production of the P11 small RNA under some conditions^3^ to control nitrogen metabolism^6^. In order to induce transgenerational (i.e., F2-F4) avoidance, *C. elegans* must ingest the small RNA (P11) made by PA14 bacteria; PA14 does not induce transgenerational avoidance when grown under all conditions, such as low temperatures and in liquid, because of this differential expression of P11^3^. In fact, this differential effect on behavior due to P11’s varying expression is what we used to identify P11 in the first place^3^. We were then able to mimic the entire P0-F4 transgenerational avoidance of PA14 effect by treating worms with *E. coli* bacteria that express the P11 sRNA. Very simply, if there is no P11 sRNA, there will be no F2-F4 avoidance. We believe that this is the crux of the issue: **the authors present no evidence that the bacteria used in their experiments produced P11 sRNA, or that the worms they tested have taken up P11 sRNA**.

## Results

### Our naïve, P0 (trained), and F2 (transgenerational) data are significant and consistent

Hunter and colleagues claim that the mild and inconsistent responsiveness of *C. elegans* to PA14 bacteria in the P0 and F1 generations they observe as support for the fact that their PA14 are active and therefore are controls for F2 behavior^1^. However, this ignores the well-established multi-factorial nature of the interactions between *C. elegans* and PA14. It is known that there are several independent paradigms that will result in P0 or F1 avoidance of PA14, but these paradigms are NOT dependent on bacterial sRNAs, and none of those transmit the information transgenerationally (F2-F4). Those paradigms include 4hr PA14 treatment, which induces avoidance in P0s via innate immunity^7^; 8 hr PA14 treatment, which causes an F1 intergenerational response but no F2 transgenerational response^8^; phenazine exposure, which induces avoidance and ASJ *daf-7p::gfp* expression in the P0 generation^9^; and intestinal bloating^10^, which induces intergenerational (F1) but no transgenerational (F2) avoidance^11^. There may well be other yet undiscovered non-sRNA-based paradigms that also induce avoidance of PA14 in the P0 and F1 generations. Thus, the P0 and F1 experiments presented here are in fact *not* positive controls for the validity of the authors’ assays, but rather may reflect these other, non-sRNA mechanisms. **Therefore, the simplest explanation for Hunter and colleagues’ inability to replicate our results is that the authors grew their PA14 bacteria under conditions that do not produce the P11 small RNA**. In these conditions we would indeed not expect them to see F2-F4 avoidance behavior; however, they might still see sRNA-independent P0 and F1 responses.

We also note that the authors even frequently fail to observe normal naïve attraction of the worms to PA14, which has been reported consistently by other labs for decades^7,12^; like other labs, we also consistently replicate this naïve attraction (**Figure 1**), again suggesting that Hunter and colleagues are not growing OP50 or PA14 correctly, or even more likely, that they have performed their choice assays incorrectly (see notes below and **Figure 4**). It should also be noted that all the wild bacteria we tested in Sengupta et al. 2024^5^, like PA14, were more attractive to the worms than lab OP50 *E. coli* under naïve conditions, pointing out again the oddness of Hunter et al.’s consistent results of OP50 *E. coli* being more attractive than PA14 to naïve worms, unlike all other groups’ findings.

**Figure 1:**
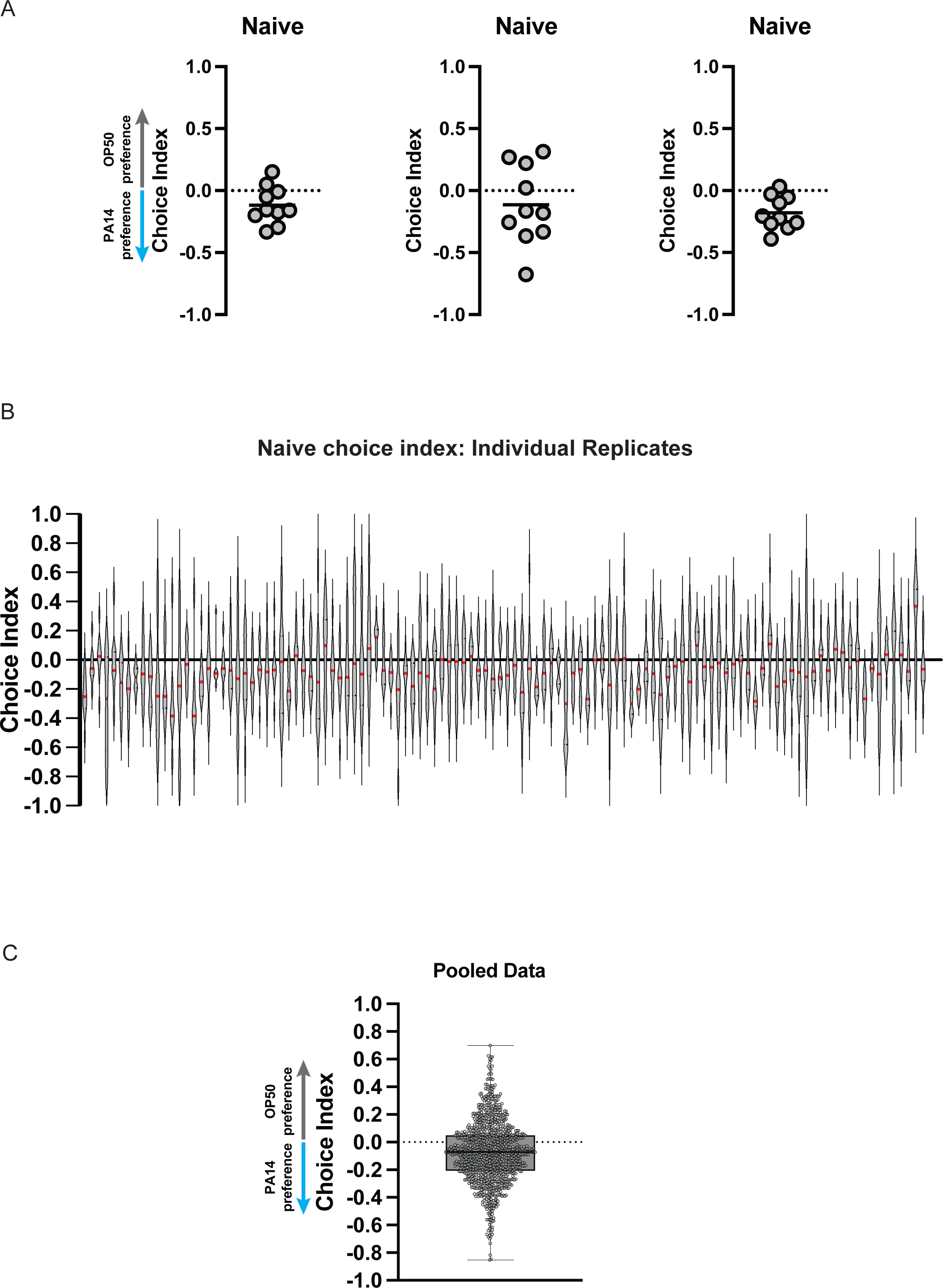
Naïve *C. elegans* reproducibly avoid PA14 in choice assays. (a) Scatter dot plots of representative replicate choice assays (OP50 versus PA14 bacteria) from naïve Day 1 adult worms (from Moore et al., 2019, Kaletsky et al., 2020, and Moore 2021). Each dot represents an individual choice assay plate (n = 10) containing ∼50-100 worms per plate (average ∼80). Choice index = (number of worms on OP50 ‐ number of worms on PA14)/(total number of worms). (b) Individual replicates of naïve choice assays from Moore et al., 2019, Kaletsky et al., 2020, and Moore 2021 are shown as violin plots. Each replicate is a separate experiment made up of 5-10 plates. The median is shown in red. (c) Individual replicates from (b) were pooled. Box plots: center line, median; box range, 25–75th percentiles; whiskers denote minimum–maximum values.

Additionally, the authors consistently fail to observe the well-established P0 avoidance response (aka olfactory learning) after training^7^. Again, in contrast, we consistently observe learned avoidance of PA14 (**Figure 2**), as other groups report^7,12^. We observe learned avoidance both from PA14 training (Figure 2, P0, blue) and from P11 small RNA training (Figure 2, P0, yellow) over hundreds of experiments. Hunter and colleagues also do not report any phenazine-induced ASJ *daf-7p::GFP* response^9^, which again suggests that the PA14 have not been grown correctly, or that their imaging assays have been performed incorrectly (see notes).

**Figure 2:**
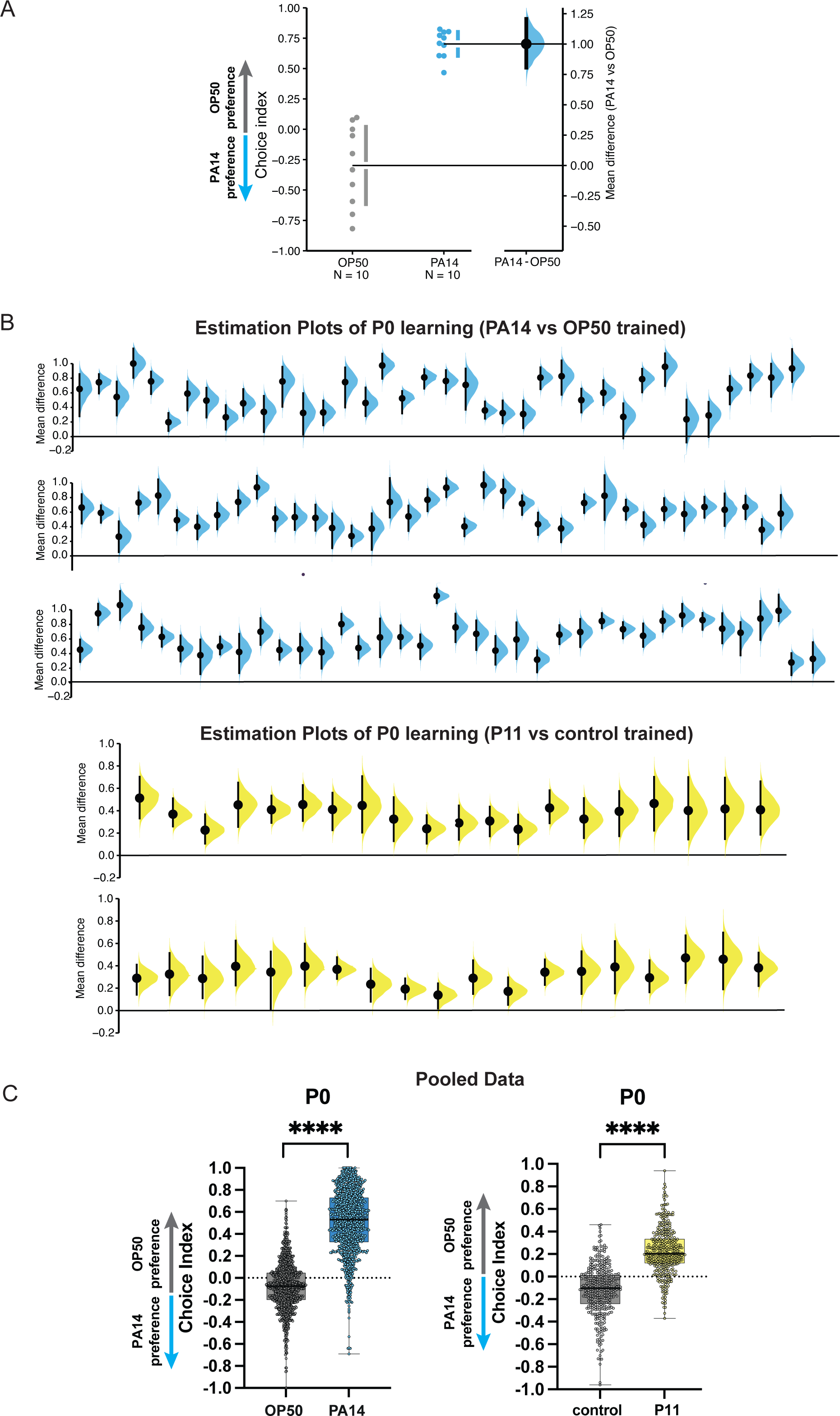
P0 worms trained on PA14 bacteria or *E. coli* expressing the P11 small RNA reproducibly avoid PA14 in choice assays. (a) A representative experiment (replicate) showing the choice index and effect size of 24h PA14-trained worms compared to the OP50-trained control. The mean difference is shown as a Gardner-Altman estimation plot. Both groups are plotted on the left axes; the mean difference is plotted on floating axes on the right as a bootstrap sampling distribution. The mean difference is depicted as a dot; the 95% confidence interval is indicated by the ends of the vertical error bar. (b) Mean differences of individual replicates from Moore et al., 2019, Kaletsky et al., 2020, and Moore 2021 are shown for PA14-vs OP50-trained mothers (blue) and P11-vs control-trained mothers (yellow). Each replicate is a separate experiment made up of 5-10 plates (average ∼80). The mean difference (effect size) is shown as a Cumming estimation plot. Each mean difference is plotted as a bootstrap sampling distribution. Mean differences are depicted as dots; 95% confidence intervals are indicated by the ends of the vertical error bars. (c) Individual replicates from (b) were pooled. Box plots: center line, median; box range, 25–75th percentiles; whiskers denote minimum–maximum values. Unpaired, two-tailed Student’s *t*-test. \*\*\*\**P* < 0.0001. Estimation graphics generated as described in Ho et al., 2019^19^.

While Hunter and colleagues claim to have followed our protocol, we observe many differences in protocols that could underlie the different results (see details below). Most importantly, they did not test the PA14 bacteria under their experimental conditions for P11 sRNA production in parallel with their experiments, which should have been the first internal control performed when there was a failure to reproduce our findings. **In summary, the authors have produced no evidence that they have used conditions that would have resulted in P11 sRNA being produced by the PA14 bacteria and/or taken up by the worms, which is consistent with their reported lack of a transgenerational effect on avoidance**. By contrast, we consistently observe avoidance of PA14 by grandprogeny (F2) of animals trained on PA14 (**Figure 3**, blue) or P11 sRNA (yellow) over dozens of experiments.

**Figure 3:**
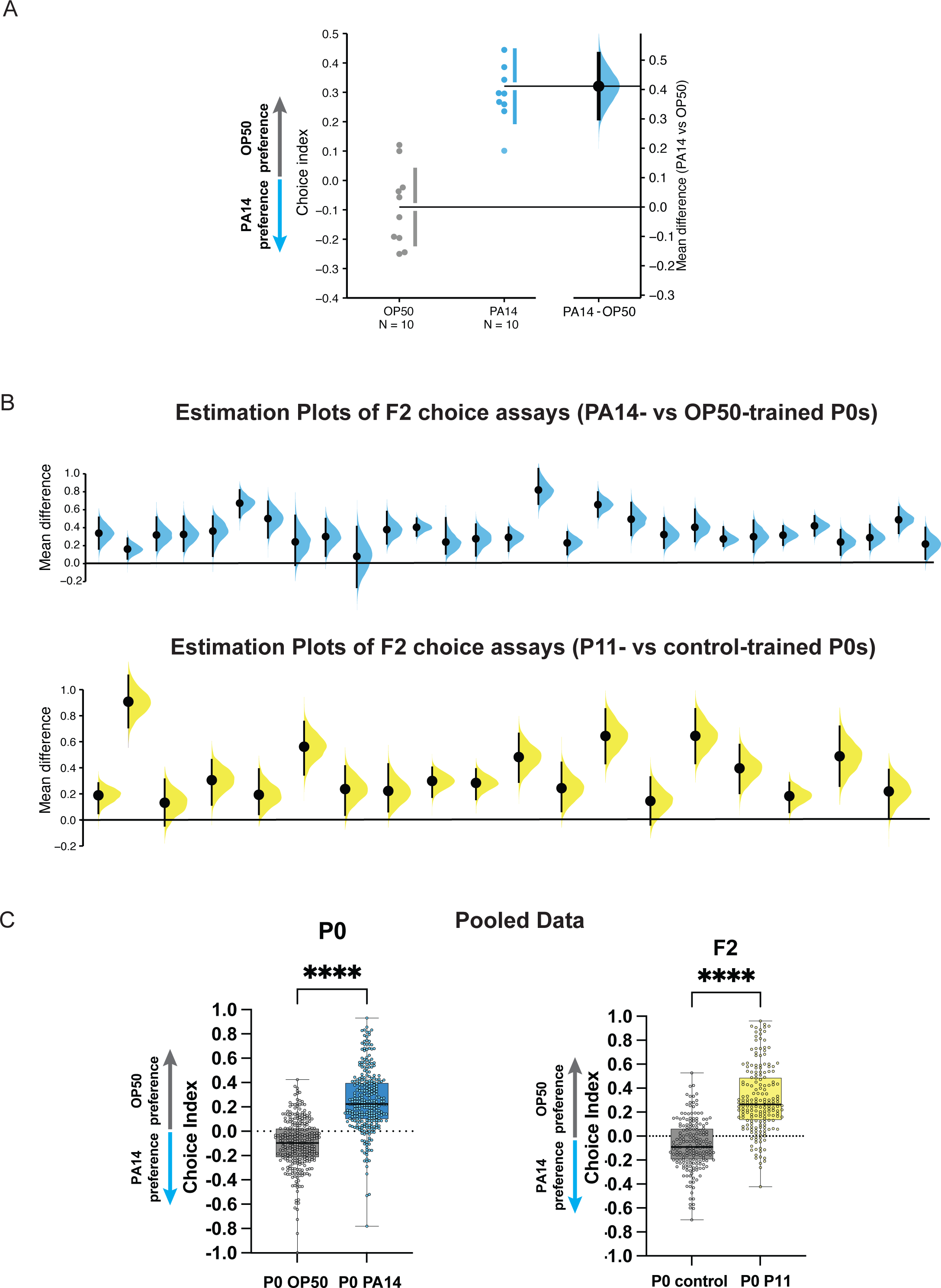
F2 progeny from P0 worms trained on PA14 bacteria or *E. coli* expressing the P11 small RNA reproducibly avoid PA14 in choice assays. (a) A representative experiment (replicate) showing the choice index and effect size of untrained (naïve) F2 animals derived from PA14-trained P0 grandmothers compared to the OP50-trained control. The mean difference is shown as a Gardner-Altman estimation plot. Both groups are plotted on the left axes; the mean difference is plotted on floating axes on the right as a bootstrap sampling distribution. The mean difference is depicted as a dot; the 95% confidence interval is indicated by the ends of the vertical error bar. (b) Individual replicates from Moore et al., 2019, Kaletsky et al., 2020, and Moore 2021 are shown for F2 worms obtained from PA14-vs OP50-trained grandmothers (top) and P11-vs control-trained grandmothers (bottom). Each replicate is a separate experiment made up of 5-10 plates (average ∼80). The mean difference (effect size) is shown as a Cumming estimation plot. Each mean difference is plotted as a bootstrap sampling distribution. Mean differences are depicted as dots; 95% confidence intervals are indicated by the ends of the vertical error bars. (c) Individual replicates from (b) were pooled. Box plots: center line, median; box range, 25–75th percentiles; whiskers denote minimum–maximum values. Unpaired, two-tailed Student’s *t*-test. \*\*\*\**P* < 0.0001. Estimation graphics generated as described in Ho et al., 2019^19^.

**Figure 4:**
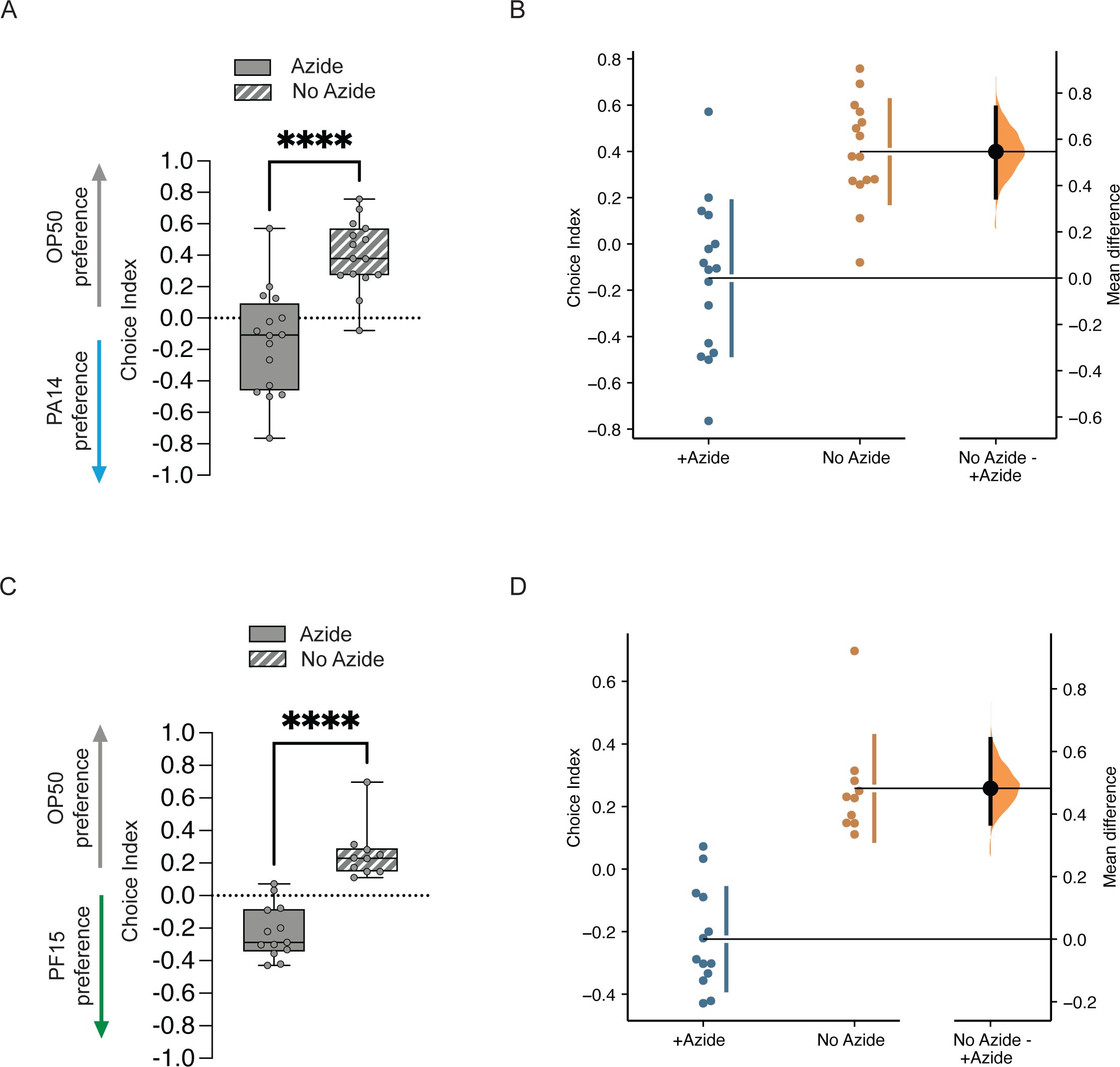
The omission of the paralytic sodium azide from choice assays has significant effects on naïve choice assays. (a) Worms were placed on choice assay plates with spots of PA14 and OP50 with or without sodium azide on the bacterial spots, then allowed to choose for one hour, followed by one hour at 4°C. No azide CI = 0.4 ± 0.06, p<0.0001. (b) Estimation plot of difference in CI between azide and no azide plates; mean difference = 0.482 [95.0%CI 0.369, 0.64]. (c) Worms were placed on choice assay plates with spots of PF15 and OP50 with or without sodium azide on the bacterial spots, then allowed to choose for one hour, followed by one hour at 4°C. No azide CI = 0.26 ± 0.05, p<0.0001. (d) Estimation plot of difference in CI between azide and no azide plates; mean difference = 0.547 [95.0%CI 0.348, 0.737].

Readers should note that in subsequent papers our lab has gone on to show that this is *not* simply a phenomenon exhibited by PA14 in lab strains of *C. elegans*: we found that wild bacteria from the worms’ habitat also induce transgenerational avoidance, and we identified the specific small RNA produced by those wild bacteria that induces the same transgenerational avoidance as we observe with PA14 and P11^5^. In contrast to the condition-specific expression of P11 in PA14, a wild *Pseudomonas* bacteria found in *C. elegans’* habitat, *P. vranovensis*^13^, seems to constitutively produce a small RNA that induces TEI avoidance, Pv1^5^. The Pv1 small RNA from *P. vranovensis* also regulates the levels of *C. elegans’ maco-1* gene through a mechanism that is similar to the P11 mechanism we previously identified. Thus, we see P0-F4 TEI avoidance induced by a wild bacteria species, *P. vranovensis*; by *P. vranovensis’* small RNAs; and by the isolated and cloned Pv1 small RNA, similar to the behavior we observed upon PA14 training under conditions that produce P11 sRNA. **That is, bacteria from a “natural setting” do indeed show “robustness” in their expression of Pv1 and transgenerational inheritance of avoidance behavior.**

Additionally, we recently found that this is an evolutionarily conserved phenomenon exhibited by yet a third *Pseudomonas* species via a third small RNA that targets yet a different *C. elegans* gene^14^. Moreover, we find that multiple wild *C. elegans* strains carry out the same behavior^4,5^. **Therefore, we are confident in the robustness and reproducibility of our work (Figures 1-3), as we have demonstrated the same sRNA-based TEI mechanism in multiple bacteria and multiple *C. elegans* strains**. The results from Sengupta et al. 2024 and Seto & Brown et al. 2024 also suggest that in fact, **the ecological significance of TEI in a natural setting in fact might be quite high**, in particular since the trained animals can also share this information horizontally through shed Cer1 capsids^4^.

Together, our data provide strong evidence that this mechanism of “reading” sRNAs from bacteria in the worm’s environment may help *C. elegans* evade microbial pathogens, as was previously proposed^15^ but was not shown until our work^2–5^. The consistency we observe over the course of our experiments in fact suggests that the transgenerational learned pathogen aversion response we identified is quite reproducible, when basic protocols are followed.

### Omission of sodium azide from choice assays may explain most differences

We were concerned about the large differences from our results that Gainey et al. reported, so we wondered whether we could identify an underlying factor that might reconcile our two labs’ findings. Gainey et al. showed differences from our results not only in their trained assays, but also in their naïve (pre-training) results. That is, their naïve CI results showed that the worms *already avoid PA14 <u>prior</u> to exposure to PA14* (CI = +0.2 to +0.4), rather than exhibiting attraction to PA14 (−0.3 to 0), as has been consistently reported by our group and others. We also noted that the Gainey et al. data lack consistency across replicates, within experiments, and across the figures of their paper, in naïve assays, P0 trained choice indices, and F1 assays (none of which depend on sRNA-induced learning).

We were particularly struck by the fact that Hunter’s group deliberately chose to not use a paralytic in their choice assays, despite the notes to do so in our protocol. Instead, Gainey et al. allowed the worms to let the worms wander for a variable amount of time on the plates with no paralytic present, then placed the worms at 4°C: “*These no-azide assay plates were moved to 4°C after 30-60 minutes*”. By contrast, the choice assay described in our TEI protocols follow standard chemotaxis assay protocols (Bargmann, *WormBook*), which use the sodium azide to paralyze the worms at the spot at their first choice. In fact, our lab carries out many chemotaxis, learning, memory, and choice assays (Kauffman et al. *PLoS Biology* 2010; Kaletsky et al. *Nature* 2016; Lakhina et al. *Neuron* 2015; Arey et al. *Neuron* 2018; Stevenson et al. *Cell Reports* 2023, etc.), and we always use sodium azide at the spot to paralyze the worms after they have made their first choice, since we are interested in measuring their immediate response to odors, rather than an adaptive response that may take place during the duration (60 min) of the choice assay itself. Similarly, we have never carried out a PA14/OP50 choice experiment without azide, since we are interested in capturing the worms’ first choice, rather than information that might change during the course of the experiment, such as adaptation or learning^16^. We wondered whether we could re-create the Hunter group’s results simply by omitting azide from the choice assay plates.

Therefore, we compared chemotaxis results after one hour of a choice assay either with or without sodium azide placed at the OP50 and PA14 spots (**Figure 4A**). Indeed, the omission of azide substantially and significantly (p<0.0001) affected the results: in the absence of a paralytic at each bacterial spot, the worms appear to flip from being to being attracted to PA14 (left, with azide, CI= - 0.15 ±0.08) as we and others have consistently shown (see **Figure 1**) to instead avoiding PA14 (right striped bar, no azide) prior to training (CI = +0.4 ± 0.06; **Figure 4A**). That is, after one hour at room temperature followed by one hour at 4°C, worms with azide stay at their initial choice of attraction to PA14, while worms on plates with no azide eventually avoid PA14, with a difference of 0.57 in CI and p<0.0001 (**Figure 4B**). These no-azide results are almost identical to the results that Gainey et al. reported for their untrained P0s, while the +azide results are similar to our previously published data, as well as published data from other labs. Furthermore, we observed the same phenomenon when we tested the effects of naïve chemotaxis to another bacteria, PF15, for which we have identified a small RNA regulator (Seto et al. 2024), that is, worms are initially attracted to PF15 as measured with azide (CI= -0.22 ± 0.05) but switch to avoidance of PF15 (0.26 ± 0.05) when allowed to wander on the plate without azide (**Figure 4C, D**). Again, the difference observed ± azide is substantial and highly significant, and would therefore obscure any attempts to measure learning induced by small RNAs.

We also observed that worms on plates transferred to the 4°C refrigerator were still moving 3 hours later, suggesting that the variable time of the assay that Hunter’s group used and placing at 4°C may have both allowed further movement of the worms well after they have migrated to their first choice. Thus, omitting azide while also carrying out choice assays for a variable amount of time and then putting the worms at 4°C, where they might continue to move, may have had significant effects both on the chemotaxis indices and on the consistency of their chemotaxis results.

We surmise that the omission of azide in the choice assay may allow the worms time to adapt or perhaps even to learn to avoid PA14 *during* the assay itself. Consistent with this hypothesis, Ooi and Prahlad (2017) previously showed that in the absence of azide, naïve worms presented with both OP50 and PA14 spots first moved towards the PA14 spot (5 min), then began to avoid PA14 and move to OP50 as the assay progressed; by 45 min, the worms have lost attraction to PA14, and by 1hr, worms have flipped from being attracted to PA14 to avoiding PA14, becoming continually more avoidant of PA14 as the assay progresses over time^16^, reminiscent of the choice assay results reported by Gainey et al. Thus, both our “no azide” results here and the Ooi & Prahlad data suggest that <u>Hunter & colleagues’ purposeful omission of azide in the choice assays caused their naïve and learned assay results to differ significantly from our published results</u>. Our results, like the Ooi & Prahlad data, are in direct contrast to Gainey et al.’s claim that “*most worms had made a choice within 15 minutes and that essentially no worm left their initial food choice during the first hour* ***(data not shown***).” Therefore, the claim by Gainey et al. that “*The addition of azide had no discernable effect on the choice assay results*” appears to be incorrect.

What are the implications of these observations? Because the magnitude of the no-azide effect on the choice assay is quite large (a shift of CI by more than 0.4), simply omitting azide from the choice assay, as Hunters’ group did, could completely mask the avoidance induced by sRNA training. That is, if the choice of untrained worms is incorrectly recorded as *already avoiding PA14*, there might be no observed avoidance due to sRNA training. (Innate immune effects in the P0 and F1 might boost the avoidance CI, but would not have any bearing on the transgenerational inheritance of avoidance due to the sRNA pathway.) Therefore, Hunter and colleagues’ decision to omit azide may explain most of the discrepancies between our results.

## Conclusions

We uncovered many experimental issues while reviewing the data from Hunter and colleagues^1^ (see **Appendix** below), but to summarize, there are enough differences between the work presented by Hunter and colleagues and ours that their data do not in fact address the validity of our findings. In fact, omission of the paralytic azide from choice plates, a deliberate deviation from our protocol, appears to be responsible for most of the differences that Gainey et al. report. Although we cannot go back in time and determine whether Gainey et al. grew PA14 under conditions that induce P11 expression or whether worm or bacterial growth conditions affected their results, we can say for certain that the omission of azide has a significant effect on the interpretation of worm training. While it is impossible to know exactly which of the many steps of our protocol that Gainey et al. changed might be the cause of their differences from our results, their decision to omit azide in their choice assays – which has major effects on the results and therefore interpretation of all subsequent data – may have led to all or most of the differences from our results that they report. By deliberately changing the choice assay protocol, it is clear that they did not try in good faith to replicate our conditions.

**More importantly, through many experiments executed by multiple authors across five publications, we have shown that transgenerational epigenetic inheritance of learned pathogen avoidance (Figure 3) is in fact robust and reliable**.

Absence of evidence is not evidence of absence: that is, the inability of Hunter and colleagues to reproduce our work, or to reproduce *any* of the previous work done on PA14 by other labs^7,9^, or to even reproduce their own results consistently within their own paper, is a flawed argument against the notion of the reproducibility of transgenerational epigenetic inheritance of pathogen avoidance in *C. elegans*.

What is important here is not the assay itself, but whether the assay can uncover meaningful biology, which it has^2–5,14^. We stand by our use of this assay, as well as the biological discoveries we have made. We are happy to invite anyone who is interested to visit our lab and learn how to carry out these protocols.

## Methods

We collected all examples of N2 naïve chemotaxis (PA14 vs OP50 choice assays) prior to training, N2 P0 and F2 choice assays after P0 training on PA14 vs OP50, and P0 and F2 choice assays after P0 training on P11 E. coli vs control E. coli from Moore et al., 2019, Kaletsky et al., 2020, and Moore 2021. These data are collected in Table S2. Multiple two-group estimation plots were generated for side-by-side comparison of multiple mean differences across independent experiments. Each mean difference is plotted as a bootstrap sampling distribution.

### Aversive learning assay

Choice plates and naïve day 1 worms were prepared as in Moore et al. 2019 and Seto et al. 2024. On the day of the assay, choice assay plates were left at room temperature for 1 h before use. For standard (sodium azide added) experiments, 1 μl of 1 M sodium azide was spotted onto each respective bacteria spot to be used as a paralyzing agent during choice assay and preserve first bacterial choice. For conditions without azide, no azide was added.

Worms were washed off training plates in 2 mL M9 into 1.5 mL tubes and allowed to pellet by gravity. Worms were washed 2-3 additional times in M9. Using a wide orifice pipet tip, 5 μl of worms (approximately 50-200 worms) were spotted at the bottom of the assay plate, midway between the bacterial spots. Plates were incubated at room temperature for 1 h before moving plates to 4C. Plates were left at 4C for 1-3h before manually counting the number of worms on each bacterial spot.

## Appendix Differences in protocols and inaccuracies in Gainey et al. 2024^1^

Hunter and colleagues used a protocol that differs significantly at many steps from our published and detailed website protocol, likely producing the extremely variable results that they report. In fact, it seems that they have not used standard bacterial growth and worm husbandry protocols, either.

When the authors contacted us, they had already developed their own protocol, which differs from ours in bacterial growth conditions, worm growth conditions, PA14 training conditions, and especially in choice assay details (see list and above, Figure 4); as they note in the text, they had already carried out most of the work presented before receiving our feedback and suggestions. The many differences in methods makes it difficult to pinpoint which might be responsible for their replication errors. We also offered to host the authors in our lab - as we have hosted others to show them the various behavioral assays in the past - but unfortunately the authors did not take us up on the offer. It appears that most of the data presented here were generated before our attempts to provide advice, and there seems to have been almost no attempt to reconcile the major protocol differences or to carry out our suggested tests and internal controls after our discussions. Outlined below are some of the general categories of differences and inconsistencies we found in their Methods. It seems deceptive to publish data generated prior to these corrections, and disingenuous to portray updates to our protocols as “deviations”. Our suggestions were made genuinely and in good faith, with the assumption that the authors wanted to use the assay, rather than using the information against us.

### 1. Inability to reproduce other groups’ naïve and trained assays

One major and surprising issue with Hunter & colleague’s results is the lack of replication of previous results that are relevant for the subsequent observation of transgenerational epigenetic inheritance of pathogen avoidance. First, they do not observe the mild attraction of naïve (untrained) animals to PA14 that is consistently reported by many other groups^7,12^ as well as our group – that is, the authors do not even observe the *initial attraction to PA14* before any training has taken place, which is well documented. Instead, their naïve chemotaxis assays are around +0.2 to +0.4 (rather than the expected -0.3 to 0, as we and others consistently observe (**Figure 1**)), suggesting that their *E. coli* OP50 are more attractive than PA14, or their PA14 is already aversive to *C. elegans* even prior to any training. This suggests that even the basic conditions of their experiments did not work, making it difficult to interpret data from post-training conditions (P0 and F1), much less transgenerational results.

**That is, in addition to being unable to replicate our transgenerational avoidance (F2) behavior, the authors show that they are unable to replicate:**

1. **Published naïve PA14 preference results**^7,12^;
2. **Published pathogen/olfactory learning (P0) results**^7,9,10^;
3. **Published *daf-7p::gfp* results**^9^;

None of these experiments depend on our lab’s protocols; they only require proper growth of PA14 and OP50, and using proper choice assays. The authors are unable to replicate the work from multiple labs, not just ours. It seems unlikely that work from the Bargmann, Kim, Zhang, Aballay and Murphy labs are all wrong; instead, the common denominator here is the present group of authors.

By contrast, we were able to consistently replicate all of the previously reported P0 and F1 findings, which is part of why we are confident in our own lab’s transgenerational (F2-F4) results.

### 2. Protocols used for choice assays

Another important difference is that the authors did not use the gold standards in the field for behavioral testing of worms (***WormBook***, *WormMethods*, Chemotaxis). Not using azide, a standardly-used paralytic, on the spot in a choice assay means that the worms will go to their first-choice spot but then they may leave, which ultimately will change the results; the worms may even be re-trained during the choice assay if not trapped at their first choice. Moving worms to a 4°C incubator, as the authors describe, will *not* immediately prevent the worms from moving away from the spot. The authors’ decision to not include azide is surprising, and likely eliminated any chance of properly capturing the worms’ first choice, or may have resulted in a different result for every assay, especially given the “30-60 minute” assay window. The authors’ reported counting method is also surprising; in our opinion, it is not feasible to accurately count more an extremely high number of worms localized in one bacterial spot; in one case the authors report that more than 700 worms were counted on a spot. Therefore, their choice assay data are unfortunately not reliable. The authors also make “*… the assumption that there are no meaningful environmental differences between choice assay plates”* – this seems like a surprising assumption to make. Unfortunately, the authors did not use the *E. coli* P11 clone we sent them to test their behavioral assay conditions. Additionally, see below for problems in treatment of the worms that may have altered behavior.

Finally, we note that the authors’ reported results for training choice indices (P0) are *extremely* variable even in wild-type worms, even in the same population, suggesting the introduction of an additional variable (or may be due to the lack of azide and thus lack of proper capture of first choice described above). Although we expect wild-type worms to exhibit some level of variability in their behavioral responses typical of wild-type bet-hedging behavior, the large variability in they report in a behavior that is usually quite stable in our hands (**Figure 2**) is unexpected. Additionally, in some cases the authors have combined learning indices from replicates that use completely different conditions (different PA14 stocks, worm growth temperatures, and light/dark conditions). Thus, they are not true replicates, and therefore, pooling them seems improper, and it is difficult to know how to interpret their data.

### 3. Lack of testing of P11 sRNA levels in their bacteria

Critically, the authors *did not check P11 sRNA levels in the PA14 grown for any of their assays*, as we suggested to them. As a result, it is not possible to know now what the levels of P11 were in any of their experiments at the time of the assays. It appears that their conditions inconsistently induce at least some but not all of the innate immune responses that have been previously reported, but they have no evidence that their bacteria produced P11 under their experimental conditions. Thus, they have not actually attempted to replicate our findings.

### 4. Bacterial growth conditions

One of the most important differences is that Hunter and colleagues use several different sets of conditions for growing their bacterial plates. Differences in PA14 growth alters the expression of P11, the small RNA that induces transgenerational (F2-F4) avoidance (which is how we originally found P11). If the authors grew the bacteria in a way that does not induce P11, such as at low temperatures or in liquid, or even on plates that were too wet, they will never see F2-F4 avoidance, even if they do see some P0 and F1 avoidance. (Also, the authors only used our PA14 bacteria in a single replicate, which is poor practice, but we doubt that the origin of PA14 is the issue here.)

Some key differences from our protocol in bacterial growth found in Gainey et al.:

a. We only treat worms on PA14 plates at 25°C; they use variable conditions.
b. Our training and choice assays, unless specified to test temperature specifically, are done at 20°C; they use variable temperatures, and even mix and match them in the same experiments.
c. We never keep seeded plates at 4°C.
d. We do not leave PA14 on plates for more than 24hrs; Prakash et al (2021) noted that PA14 produces repulsive odors (1-undecene) after 24hrs^17^, so old PA14 plates may have caused some of the high naïve avoidance the authors report.
e. We don’t treat worms on PA14 at 15°C, as the authors mention; if at some point the worm was on a plate of PA14 at 15°C the authors are only studying the *innate immune* part of the pathway, because at 15°C there is no P11 expression^3^, and thus there could be no transgenerational effect.
f. We only use fresh OP50, not “stored for up to 1 month”. (We note that the bioRxiv version of this description was changed from a version we were sent just two weeks ago.)
g. Bacterial plates should never be wet, which might change their growth conditions and alter P11 expression.

### 5. Harsh training/growth conditions

The authors’ training conditions seem to be extremely stressful for the animals, as they report that “15-30% of the PA14 cultured worms were found desiccated on the walls of the plate” – we have never seen this extreme level of death or escape in our assays in the 24hrs of training, nor have we seen such low numbers of progeny produced, or seen delays in their development. This corroborates that the plates or bacterial growth conditions - essential components of these assays - are notably different. The inconsistency of their P0 *daf-7p::gfp* imaging results (and deviation from all other labs’ results), where they observe no signal in P0 but see it in F1, also suggests that the worms are very sick after training. (Perhaps differences in start of training as a result of overbleaching-induced developmental delays also contribute to these problems, we are not sure.) In any case, the worms themselves appear to be very sick in the P0 generation, which may have led to their extreme censoring, low progeny numbers, delays in development, and changes in behavior.

Interestingly, extremely stressful conditions, such as starvation, can “reset” or erase transgenerational effects^18^ – it is possible that if their worms are very sick and stressed, as they seem to be, it may have affected the inheritance, but that presumes that they were even exposed to P11.

### 6. Harsh treatment of worms that can affect behavior

We do not know how important the many other changes the authors made to the protocol are, but since our lab has carried out learning and memory behavior assays for almost 20 years, there are specific common procedures that we never do to worms in any of our lab’s behavioral assays that we note are present in their Methods, but may severely stress the worms and thus interfere with behavior.

We note that the authors added these steps to our protocol that we do not use on worms that will be tested for behavior:

a. We never pellet or vortex worms, which can physically damage them and their neurons.
b. We never use Triton X100, which may permeabilize eggs;
c. Overbleaching: Their bleaching protocol is very harsh on the eggs (2x bleaching, totally dissolving mothers) – in our lab, we bleach just long enough to release the eggs. Overbleaching eggs can stress the worms, delay development, and affect behavior in adulthood.

### 7. Procotols used for imaging

The authors’ reported *daf-7p::gfp* intensities do not correlate with the position of the neuron within the animal (in some cases, the neuron farther away from the objective is brighter); by averaging, the authors may have averaged out biologically relevant differences. Other differences with our imaging conditions include sealing the slides, which may affect aeration and the duration of *daf-7p::gfp* expression in live animals; we also carry out imaging as quickly as possible. The authors also excluded data, including high or low *daf-7p::gfp* intensities, for no apparent reason, even though it is obvious that there can be true extreme biological differences in *daf-7p::gfp* expression in individual worms. (Stressing the worms might have some effects on *daf-7* expression or GFP fluorescence.)

However, given the issues with bacterial growth issues described above, we do not think that simple imaging method differences can account for the issues.

### 8. General factual errors

In addition to these differences, the authors have misrepresented the number of worms used in our papers. (*“*.. *details in this and subsequent reports suggest that their sample sizes in some choice assays may have been unreasonably small (as few as 10-20 worms per assay plate) (Kaletsky et al*., *2020; Moore et al*., *2021b)*”).

In fact, as shown in the table below, we generally **use ∼50-90 worms/plate, with a mean of 86 and median of 75 worms/plate**. In every experimental replicate **we use ∼8 choice assay plates per training condition, and multiple (∼3) replicates for any experiment**. Therefore, even if there were a few outlier plates with low numbers in a single replicate in a single experiment, it would have little effect on our conclusions.

|  | Mean worms/plate | Median worms/plate | Total assay plates | Total worms |
| --- | --- | --- | --- | --- |
| Moore et al., 2019 | 106.5 | 93 | 1447 | 154163 |
| Kaletsy et al., 2020 | 89.2 | 76 | 11085 | 988699 |
| Moore et al., 2021 | 84.1 | 75 | 15646 | 1315348 |
| Sengupta et al., 2024 | 76.3 | 66 | 3940 | 300684 |
| Combined papers | 85.9 | 75 | 32118 | 2758894 |

To summarize, our findings are both consistent and significant. Therefore, the authors’ implication that we have used low n’s of worms to interpret any of our results is simply incorrect and is a gross misrepresentation of our work.

However, it is important to note that we did not *need* to analyze this large number of worms and plates because of assay “variability” or to achieve data significance to make our observations, as the changes in behavior we observe are quite obvious in most cases. In fact, it should be remembered that we found the transgenerational inheritance of PA14 avoidance serendipitously in the course of carrying out studies of the effects of pathogenesis on developmental delays (Moore et al, unpublished), specifically because the transgenerational effect was so obvious.

### 9. “Personal communication”

It should be noted that we did not approve of any of the uses of “Personal Communication” cited in this article, which misconstrue our repeated efforts to try to help the Hunter lab carry out their work. We are happy to provide emails regarding this point.

## Notes

### Competing Interest Statement

The authors have declared no competing interest.

### Summary of Updates

We have added new data testing the conditions that Hunter and colleagues used in their choice assays, where they omitted the paralytic sodium azide from the assays altogether, claiming there was no effect. We find that there is a substantial and significant effect in both PA14 and PF15 choice assays (Figure 4), and the effect is so large that this omission would have changed the results enough to mask any effects from small RNA-mediated training. The authors also wrote about purposely omitting sodium azide, even though we specified the requirement for sodium azide in our protocol - thus nullifying their claim that they followed our protocol. Therefore, Hunter and colleagues' claims that our results are "irreproducible" are shown to be false by these data.

## References

1. Gainey, D.P., Shubin, A.V., and Hunter, C.P. (2024). Irreproducibility of transgenerational learned pathogen-aversion response in C. elegans. Preprint, https://doi.org/10.1101/2024.06.01.596941 10.1101/2024.06.01.596941.

2. Moore, R.S., Kaletsky, R., and Murphy, C.T. (2019). Piwi/PRG-1 Argonaute and TGF-β Mediate Transgenerational Learned Pathogenic Avoidance. Cell 177, 1827-1841.e12. 10.1016/j.cell.2019.05.024.

3. Kaletsky, R., Moore, R.S., Vrla, G.D., Parsons, L.R., Gitai, Z., and Murphy, C.T. (2020). C. elegans interprets bacterial non-coding RNAs to learn pathogenic avoidance. Nature 586, 445–451. 10.1038/s41586-020-2699-5.

4. Moore, R.S., Kaletsky, R., Lesnik, C., Cota, V., Blackman, E., Parsons, L.R., Gitai, Z., and Murphy, C.T. (2021). The role of the Cer1 transposon in horizontal transfer of transgenerational memory. Cell 184, 4697-4712.e18. 10.1016/j.cell.2021.07.022.

5. Sengupta, T., St. Ange, J., Kaletsky, R., Moore, R.S., Seto, R.J., Marogi, J., Myhrvold, C., Gitai, Z., and Murphy, C.T. (2024). A natural bacterial pathogen of C. elegans uses a small RNA to induce transgenerational inheritance of learned avoidance. PLOS Genet. 20, e1011178. 10.1371/journal.pgen.1011178.

6. Marogi, J.G., Murphy, C.T., Myhrvold, C., and Gitai, Z. (2023). P. aeruginosa controls both C. elegans attraction and pathogenesis by regulating nitrogen assimilation. Preprint, https://doi.org/10.1101/2023.11.29.569279 10.1101/2023.11.29.569279.

7. Zhang, Y., Lu, H., and Bargmann, C.I. (2005). Pathogenic bacteria induce aversive olfactory learning in Caenorhabditis elegans. Nature 438, 179–184. 10.1038/nature04216.

8. Pereira, A.G., Gracida, X., Kagias, K., and Zhang, Y. (2020). C. elegans aversive olfactory learning generates diverse intergenerational effects. J. Neurogenet. 34, 378–388. 10.1080/01677063.2020.1819265.

9. Meisel, J.D., Panda, O., Mahanti, P., Schroeder, F.C., and Kim, D.H. (2014). Chemosensation of Bacterial Secondary Metabolites Modulates Neuroendocrine Signaling and Behavior of C. elegans. Cell 159, 267–280. 10.1016/j.cell.2014.09.011.

10. Singh, J., and Aballay, A. (2019). Intestinal infection regulates behavior and learning via neuroendocrine signaling. eLife 8, e50033. 10.7554/eLife.50033.

11. Hong, C., Lalsiamthara, J., Ren, J., Sang, Y., and Aballay, A. (2021). Microbial colonization induces histone acetylation critical for inherited gut-germline-neural signaling. PLOS Biol. 19, e3001169. 10.1371/journal.pbio.3001169.

12. Ha, H., Hendricks, M., Shen, Y., Gabel, C.V., Fang-Yen, C., Qin, Y., Colón-Ramos, D., Shen, K., Samuel, A.D.T., and Zhang, Y. (2010). Functional Organization of a Neural Network for Aversive Olfactory Learning in Caenorhabditis elegans. Neuron 68, 1173–1186. 10.1016/j.neuron.2010.11.025.

13. Samuel, B.S., Rowedder, H., Braendle, C., Félix, M.-A., and Ruvkun, G. (2016). Caenorhabditis elegans responses to bacteria from its natural habitats. Proc. Natl. Acad. Sci. 113. 10.1073/pnas.1607183113.

14. Seto, R.J., Brown, Rachel, Kaletsky, R., Parsons, L.R., Moore, R.S., and Murphy, C.T. (2024). Pseudomonas fluorescens 15 small RNA Pfs1 mediates transgenerational epigenetic inheritance of pathogen avoidance in C. elegans through the Ephrin receptor VAB-1. Preprint.

15. Winston, W.M., Molodowitch, C., and Hunter, C.P. (2002). Systemic RNAi in C. elegans Requires the Putative Transmembrane Protein SID-1. Science 295, 2456–2459. 10.1126/science.1068836.

16. Ooi, F.K., and Prahlad, V. (2017). Olfactory experience primes the heat shock transcription factor HSF-1 to enhance the expression of molecular chaperones in C. elegans. Sci. Signal. 10, eaan4893. 10.1126/scisignal.aan4893.

17. Prakash, D., Ms, A., Radhika, B., Venkatesan, R., Chalasani, S.H., and Singh, V. (2021). 1-Undecene from Pseudomonas aeruginosa is an olfactory signal for flight-or-fight response in Caenorhabditis elegans. EMBO J. 40, e106938. 10.15252/embj.2020106938.

18. Houri-Zeevi, L., Teichman, G., Gingold, H., and Rechavi, O. (2021). Stress resets ancestral heritable small RNA responses. eLife 10, e65797. 10.7554/eLife.65797.

19. Ho, J., Tumkaya, T., Aryal, S., Choi, H., and Claridge-Chang, A. (2019). Moving beyond P values: data analysis with estimation graphics. Nat. Methods 16, 565–566. 10.1038/s41592-019-0470-3.

